# Gut microbiota phospholipids regulate intestinal gene expression and can counteract the effects of antibiotic treatment

**DOI:** 10.1101/2025.01.28.633906

**Authors:** Xue-Song Zhang, Yujue Wang, Haipeng Sun, Christa Zerbe, Emilia Falcone, Sneha Bhattacharya, Meifan Zhang, Zhan Gao, Maria Elena Diaz-Rubio, Disha Bharj, Disha Patel, Samuel Pan, Gabrielle Ro, Jessica Grenard, Abigail Armstrong, Yue Sandra Yin, Maria Gloria Dominguez-Bello, Steven Holland, Xiaoyang Su, Martin J. Blaser

## Abstract

The gut microbiome influences immune and metabolic homeostasis. Our research using non-obese diabetic (NOD) mice revealed that early-life antibiotic exposure remodels the gut microbiome affecting metabolism and accelerating type 1 diabetes (T1D) incidence, with cecal material transplant (CMT) mitigating the damage. Now examining murine intestinal lipidomic profiles, we identified 747 compounds. Comparing the lipidomic profiles of cecal contents of conventional and germ-free mice and their diets, we identified 87 microbially-produced lipids reduced by antibiotic exposure but CMT-restored. Parallel analysis of human fecal lipid profiles after azithromycin-exposure showed significant alterations with substantial overlap with mice. *In vitro* co-culture with mouse macrophages or small intestinal epithelial cells and human colonic epithelial cells identified phospholipids that repress inflammation through the NF*κ*B pathway. Oral administration of these phospholipids to antibiotic-treated NOD mice reduced expression of ileal genes involved in early stages of T1D pathogenesis. These findings indicate potential therapeutic anti-inflammatory roles of microbially-produced lipids.

## Introduction

Gut microbes synthesize diverse small molecules that significantly influence host metabolism, immunity and neural systems affecting health and disease^1-3^. Early life is the critical period for establishing the gut microbiome and facilitating healthy host development^4-6^. Environmental factors altering gut microbiota composition can increase risk of subsequent metabolic and immune-mediated diseases^7,8^. Type 1 diabetes (T1D), an autoimmune disease usually beginning in childhood, may be influenced by gut microbiota-host interactions^9-12^. Our recent studies using non-obese diabetic (NOD) mouse models showed that early-life antibiotic exposure perturbs the gut microbiota, interferes with innate and adaptive immune effectors, alters ileal gene developmental patterns, and increases T1D onset incidence^13-15^. Maternal cecal microbiota transplant (CMT) to NOD mice after early-life antibiotic exposure significantly mitigated the induced T1D enhancement by partially restoring microbial diversity, abundance of specific taxa and altered metabolic pathways, including these related to bacterial lipid/fatty acid metabolism^13-15^. Here we comprehensively investigated the profiles of gut microbially-produced lipid molecules, evaluating the effects of early-life antibiotic perturbation and CMT on lipid profiles. We also extended our research to humans by examining the effects of administering oral antibiotics on microbial lipid profiles of fecal specimens obtained from young adults enrolled in the NIH-MIME study (Microbial, Immune and Metabolic Interference by Antibiotics)^16^. We identified specific microbially-produced lipids that affect gut innate immune factors *in vitro* and *in vivo*. These studies indicate the therapeutic potential of these bacterial lipids for preventing/ treating gut inflammatory and immune-associated diseases.

## Results

### Cecal microbiota lipid metabolism pathways in NOD mice were perturbed by early life antibiotic exposure and partially restored by cecal material transplant

In our prior studies of NOD mice, a single early-life antibiotic course (1P) perturbed gut microbiota, increasing T1D incidence compared to control (C), while CMT partially restored microbiota and reduced the increased risk^15^. Metagenomic analysis of fecal microbiomes identified 1501 bacterial metabolism pathways, indicating significant differences between the three treatment groups (**Figure 1A**)^15^. We now extend the analysis to the 36 bacterial lipid metabolism pathways affected by 1P but restored by CMT (**Figure 1B**). Unsupervised clustering of the 23 high-prevalence pathways segregated C and 1P samples, while CMT samples mostly clustered with C (**Figure 1C**), consistent with recovery from the accelerated T1D^15^.

**Figure 1.**
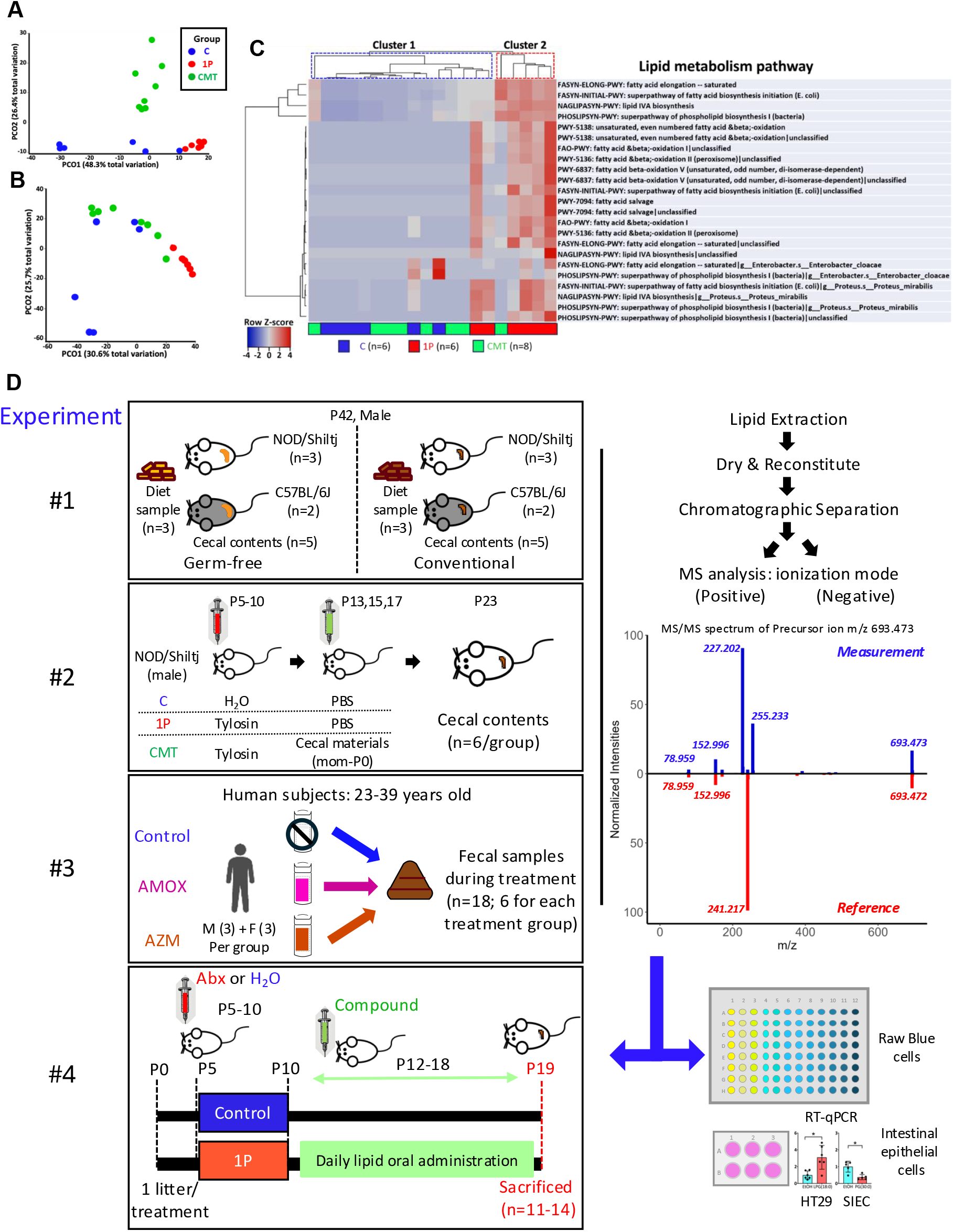
Effect of early-life antibiotic treatment and cecal material transplant (CMT) on fecal bacterial lipid metabolism pathways in mouse feces and subsequent design of lipidomic studies. Analyses based on fecal metagenomic sequencing of P23 male NOD mice in three treatment groups (C, Control; 1P, 1PAT; CMT, CMT after 1PAT). Bray-Curtis distances for pathways identified in the metagenomic analysis with pair-wise PERMANOVA analysis: **A)** 1501 metabolic pathways; **B)** 36 lipid metabolic pathways; **C)** Heatmap of the 23 identified high-prevalence lipid metabolism pathways. In unsupervised hierarchical clustering, the 1P samples were significantly overrepresented in Cluster 2 (p=0.0139; FET). **D)** Study designs for lipidomic analyses and for administering defined lipid compounds to NOD mice. **Experiment 1:** Cecal contents of 6-week-old C57BL/6J and NOD/ShiLtJ GF and conventional mice and their dietary materials. **Experiment 2:** Cecal contents from male NOD mice at P23 in C, 1P, CMT mouse-groups (n=6/group). **Experiment 3:** Fecal samples from human volunteers during treatment in the MIME Study, who received one week of (i), azithromycin (AZM), (ii), amoxicillin (AMX), or (iii), no antibiotic (Control). **Experiment 4**: *In vitro* co-culture with macrophage Raw blue cells to assay immune key regulator NF*κ*B level with colorimetric reaction and with intestinal epithelial cells (colonic HT29 cells & mouse small intestinal epithelial cells) to assay immune gene expression via RT-qPCR, followed by *in vivo* experiment with early-life antibiotic exposure followed by restorative treatment. Pregnant NOD/ShiLtJ mice were randomized into 8 groups and pups studied: 7 groups received tylosin (1PAT; 1P) during days P5-P10. From P12-P18, mice in five 1P groups were each gavaged daily with a defined lipid [LPG(13:0), PG(15:0_15:0), LPG(16:0), LPG(18:0), or retinoic acid [RA, positive control], or with CMT from healthy male P23 NOD pups, or solvent (PBS-2% EtOH) only. Another group was gavaged with solvent from P12-P18 as a control. From each group, all male pups from two litters were sacrificed at P19 to assess effects of the administered lipid on intestinal gene expression.

### Gut microbiome lipidomic and diet lipidomic analysis in Germ-free and conventional mice

To determine the origins of gut lipids, we conducted lipidomic analyses of both diets and cecal contents of germ-free (GF) and conventional mice (**Figure 1D, top panel**). MS/MS analysis identified 747 non-redundant lipid compounds including fatty acids, mono-, di-, and triacylglycerols, ceramides, multiple phospholipids, sterol esters sulfonolipids, sphingolipids, cardiolipins, and carotenoids (Full list: **Table S1**). Of the 747 lipids, 230 and 517 had odd-chain or even-chain fatty acid structures, respectively; odd-chain lipids are nearly universally of bacterial origin, whereas even-chain lipids may be of either bacterial or eukaryotic origin^17^.

Pair-wise PERMANOVA analysis indicated that: lipid profiles of cecal contents from the conventional and GF mice were distinct from the diets; significant differences between conventional and GF mice; and the two mouse strains had similar profiles according to condition (**Figure 2A**). Unsupervised hierarchical clustering analysis revealed that all diet samples clustered distinct from all mouse samples and that all GF mice clustered together and was separated from the cluster of the conventional mice (**Figure 2B**). We identified a group of 349 dietary lipids that were highly absent in the cecal contents, likely metabolized by the host proximal to the cecum (**Figure 2B, green box**). There was a group of 94 lipids low in diets, but abundant in all mice regardless of whether a microbiome was present; we considered these host-produced lipids (**Figure 2B, yellow box**). We identified 144 lipid compounds, abundant in conventional cecal contents from conventional mice but not in germ-free or in diets (**Figure 2B, blue boxes**); we considered these as gut microbially produced lipids (GMPLs), which were the focus of further studies. Conventional and GF mice were more distinct in their odd-chain (**Figure 2C**) than even-chain (**Figure 2D**) lipids, consistent with a higher proportion of microbially-produced odd-chain than even-chain lipids. Unsupervised hierarchical clustering also showed a higher proportion of microbially-produced lipid compounds within the odd-chain compounds (**Figure S1**). Using criteria based on statistical significance (adjusted p<0.05), and fold-differences (FD), we identified 59 odd-chain and 28 even-chain GMPLs, dominated by phospholipids and sphingolipids (**Figure 2EF**).

**Figure 2.**
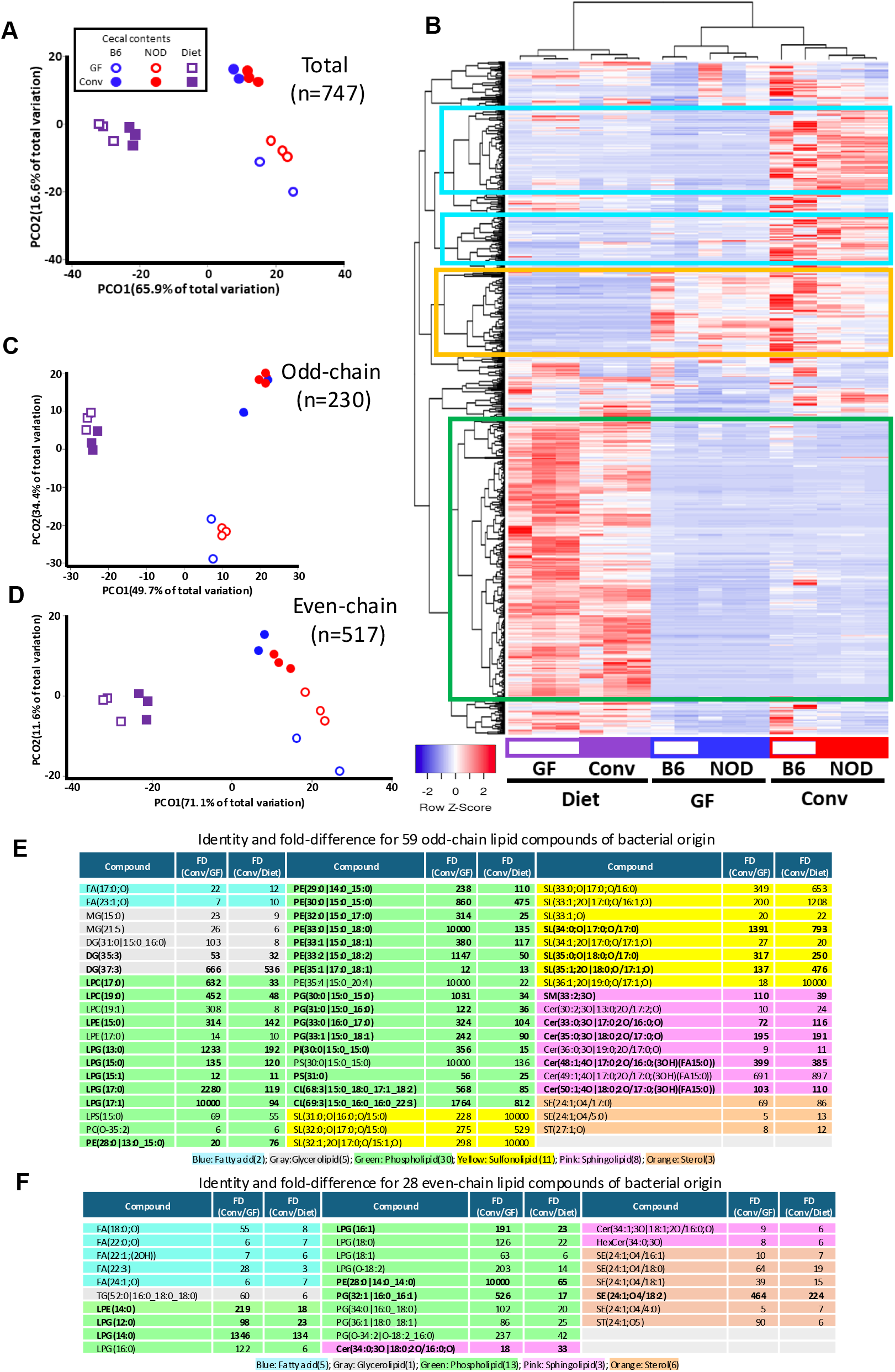
Lipidomic analysis of cecal contents of 6-week-old Germ-free (GF) and conventional mice and their diets (Experiment 1). **A)** Bray-Curtis distances of the 747 identified lipids. Differences between conventional vs. GF mice; conventional mice vs. conventional diet; GF mice vs GF diet. Adjusted p values were <0.02 for each comparison, by pairwise PERMANOVA analysis. **B)** Unsupervised hierarchical clustering based on 747 lipid compounds. Green box: lipids present in the diets metabolized by hosts regardless of microbiota presence or absence. Yellow box: mouse-produced lipids, present in both conventional and GF hosts. Blue boxes: microbially-produced lipids, abundant in conventional cecal contents but not in diets or in cecal contents from GF mice. Bray-Curtis distance matrix: **C)** 230 odd-chain; **D)** 517 even-chain microbially-produced lipid compounds. For both panels, pair-wise PERMANOVA analysis indicated significant differences (adjusted p <0.02) for the same comparisons as in panel A. Identification of 59 significant microbially-produced lipid compounds: **E)** 59 odd-chain; **F)** 28 even-chain. Defined by FD(_Conv/GF)_ & FD_(Conv/Diet)_ >5; compounds meeting both FD-criteria and statistical significance (T-test p<0.05) in bold.

We further conducted lipidomic analysis using six cultured gut bacterial strains representing different phyla—*Alistipes finegoldii* (Bacteroidota), *Muribaculaceae* (Bacteroidota), *Bifidobacterium longum* (Actinomycetota), *Escherichia coli* (Pseudomonadota), *Clostridium perfringens* (Bacillota), and *Lactobacillus murinus* (Bacillota). Bray-Curtis distance analysis based on the 391 lipid compounds identified (**Table S2**) revealed distinct lipid profiles for each (**Figure S2A**), as did unsupervised clustering (**Figure S2B**). This analysis provides evidence that gut bacterial taxa contribute distinctively to the overall lipid diversity within the microbiome.

### Effects of early-life antibiotic exposure on lipid profiles of NOD mouse cecal contents

Next, we investigated the effects of early-life antibiotic exposure GMPLs in cecal contents of C, 1P and CMT NOD mice^15^ (**Figure 1D**). Using standardized methods for the 747 lipid compounds (**Table S1**), we found that 1P mice had significantly altered profiles compared to C, while CMT restored the profile toward control, as determined by Bray-Curtis distance analysis with pair-wise PERMANOVA (**Figure 3A**). Unsupervised clustering revealed two distinct subgroups: one with all six 1P and two of the six CMT samples, while the other with all six control and four of the six CMT samples (**Figure 3D**). This dichotomy indicates that 1P significantly remodeled the cecal lipid profiles with partial restoration by CMT. A group of 343 (45.9%) were decreased by antibiotics and restored by CMT (**Figure 3D, blue box**), whereas 99 (13.3%) were increased by antibiotics and restored by CMT (**Figure 3D, red box**); 1P significantly altered both odd-chain and even-chain profiles compared to control (**Figure 3BC**), and CMT restored profiles toward control (**Figure 3BC**). We identified 49 GMPLs affected by the 1P treatment (**Figure 3E**). Further analysis of the 1P effects on the 747 compounds and the three subgroups defined in **Figure 2B**, GMPLs (blue boxes), host-produced compounds (yellow box), and diet-derived compounds (green), using fold-difference and statistical significance criteria confirmed the extent of antibiotic impact on lipid profiles, particularly GMPLs (**Figures 3F, S3**).

**Figure 3.**
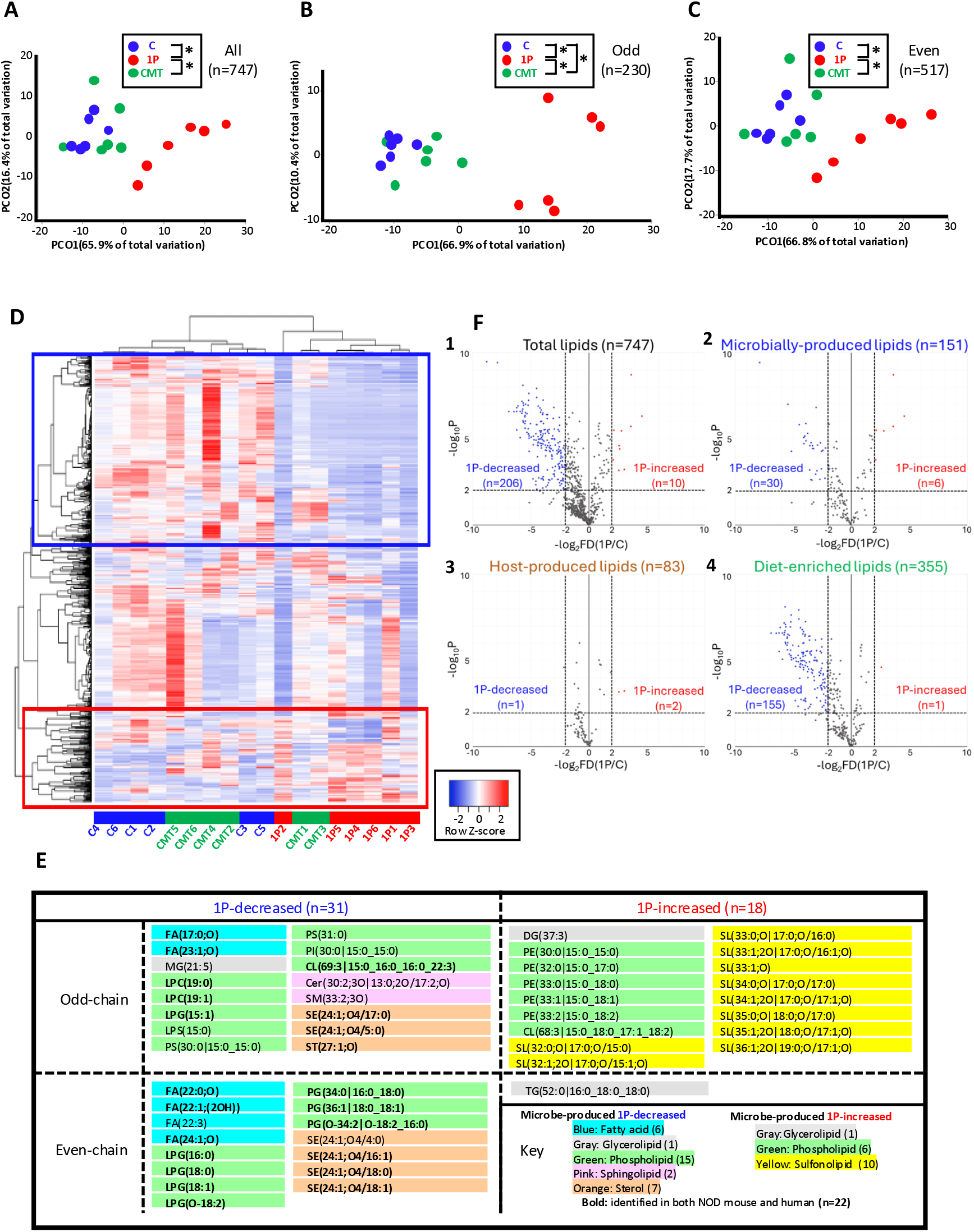
Antibiotic effects on cecal content lipid profiles in NOD mice (Experiment 2). **A)** Bray-Curtis distances for the 747 identified mouse cecal compounds. Significant differences between C & 1P and between 1P & CMT (adjusted p-value =0.012 for both, pairwise PERMANOVA). **B)** Bray-Curtis distances for the 230 identified odd-chain lipid compounds. Significant differences [pairwise PERMANOVA analysis] between C & 1P (adjusted p-value =0.003), between 1P & CMT (adjusted p-value =0.006), and between C & CMT (adjusted p-value =0.024). **C)** Bray-Curtis distances for the 517 even-chain identified compounds. Significant differences [pairwise PERMANOVA analysis] between C & 1P (adjusted p-value =0.006), between 1P & CMT (adjusted p-value =0.012). No significance between C and CMT (adjusted p-value =0.30). **D)** Unsupervised clustering based on the 747 compounds. Blue box: i lipid compounds decreased by 1P (compared with control) and restored by CMT (n=343; 45.9%). Red box: lipid compounds increased by 1P and restored by CMT (n=99, 13.3%). **E)** Microbially-produced lipids with 1P altering abundance. **F)** Volcano plots showing the impact of tylosin treatment on the abundances of the compounds in the Control and 1P groups. Panels: 1) All 747 compounds; 2) 151 microbially-produced compounds identified from **Figure 2B** [blue boxes]; 3) 83 host-produced compounds [yellow box]; 4) 355 diet-derived compounds [green box]. Blue points represent 1P-decreased compounds, with geometric mean fold-difference >4 and log-transformed p-value <0.01. Red points represent 1P-increased compounds, using the same criteria.

### Effects of azithromycin on fecal lipid profiles of human subjects

In parallel, we analyzed the fecal microbiome and lipidomics of young adult human subjects (**Figure 1D)**, who received azithromycin (AZM), amoxicillin (AMX), or no antibiotic (Control). At the 16S level, azithromycin did not impact α-diversity (**Figure 4A**), but did alter β-diversity (**Figure 4B**). We focused on the azithromycin-perturbed lipids. MS/MS analysis identified 728 lipid compounds (228 odd-chain and 500 even-chain compounds) in the human fecal contents (**Table S1**) lacking only 17 phospholipids and 2 sphingolipids from mouse cecal contents. Bray-Curtis distances were significant between the control and azithromycin group, but not for the amoxicillin group, consistent with the 16S studies (**Figure 4C**). Unsupervised clustering of the 18 subjects from the 3 groups based on the 728 lipids showed two distinct clusters: Cluster 1 includes 5 of the 6 AZM-exposed subjects and 1 AMX-exposed subject. Cluster 2 includes all 6 control subjects, 4 of the 6 AMX-exposed subjects and 1 AZM-exposed subject (**Figure S4**). Among the 59 odd-chain GMLs (**Figure 2E**), azithromycin significantly decreased 31 (53%) and increased none. Of the 28 even-chain GMLs (**Figure 2F**), azithromycin significantly decreased 20 (71%) and increased none. Thus, the lipids affected by the macrolide-treatments in the mouse and human intestinal samples significantly overlapped (**Figure 4D**). Volcano plot analysis (**Figure 4E**) confirmed the significant impact of azithromycin on microbially produced lipids, identifying 39 compounds that were decreased, with none increased (**Figures 4E, S5**; amoxicillin treatment significantly altered only 3 lipids (**Figure S5**).

**Figure 4.**
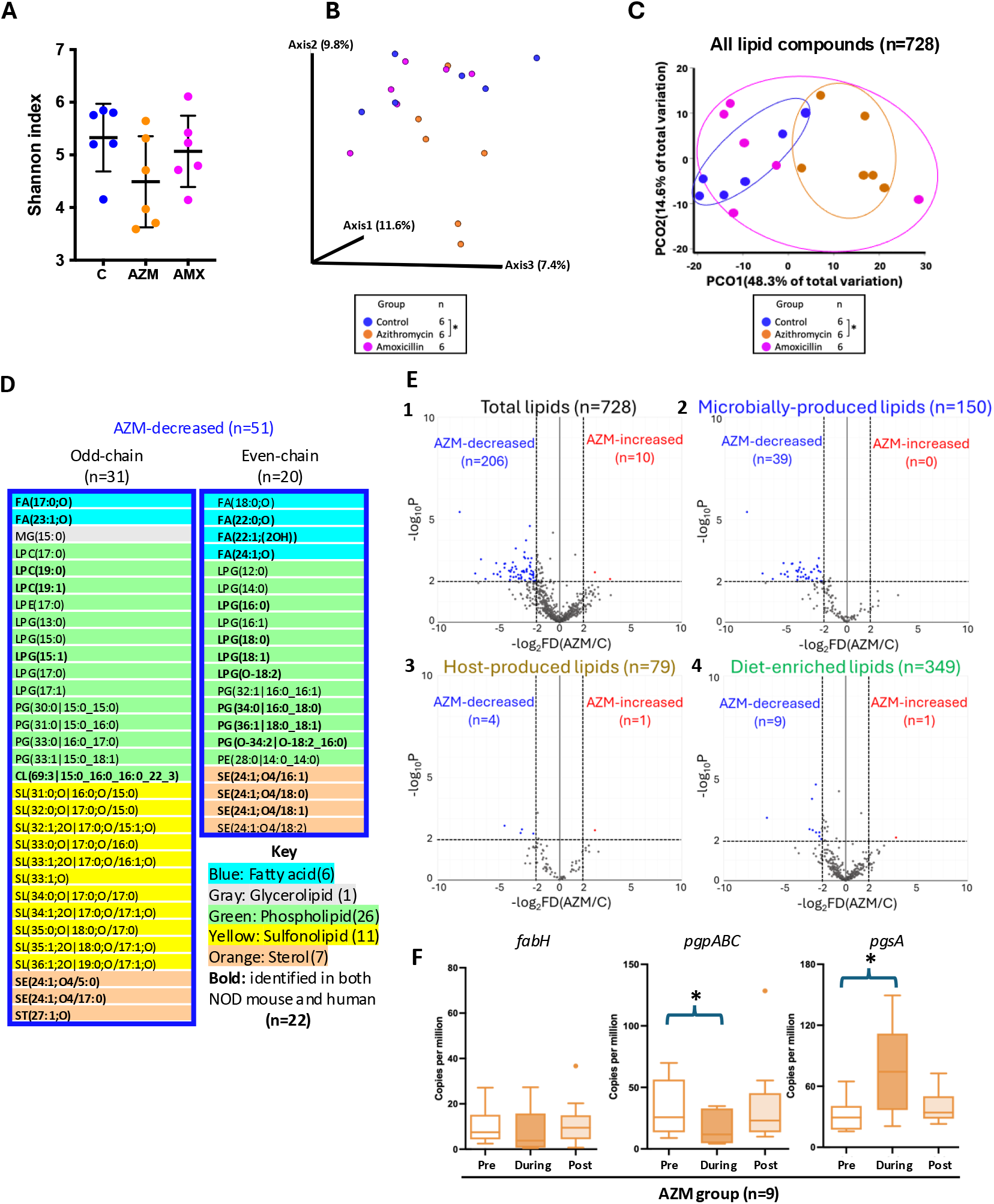
Antibiotic effects on fecal microbiome and lipid profiles in human volunteers (Experiment 3). **A)** Fecal α-diversity: Shannon index, depth 14,246 reads. No significant differences observed (Control vs. azithromycin groups p=0.078). **B)** Fecal β-diversity: Jaccard distance matrix showing significant intergroup distances (*p=0.02; pairwise PERMANOVA), depth 14,246 reads. **C)** Bray-Curtis distances for the 728 lipid compounds identified in the human feces, including 228 odd-chain and 500 even-chain compounds; pair-wise PERMANOVA analysis indicated significant differences between the control and azithromycin groups (* adjusted p-value=0.015). **D)** Microbially-produced lipids differentiated by azithromycin include 31 odd-chain (all decreased) and 20 even-chain (all decreased) compounds; no compounds were significantly increased by azithromycin. The 22 lipids significantly affected in both antibiotic-treated mice and humans are shown in bold. **E)** Volcano plots showing the impact of azithromycin treatment on the abundances of the nonredundant lipid compounds. Panels: 1) All 728 identified compounds, 2) 150 microbially-produced compounds identified from **Figure 2B** blue boxes; 3) 79 host-produced compounds from yellow box; 4) 349 diet-derived lipid compounds from green box. **F)** Fecal metagenomics: normalized number of copies of the genes encoding the key enzymes involved in bacterial (Lyso-)PG biosynthesis before, during, and after the azithromycin treatment. *p<0.05 compared with 1P group; Mann-Whitney *U*-test.

Phospholipids were the largest group of GMPLs decreased by azithromycin (51%) (**Figure 4D**), similar to the findings in mice (**Figure 3E**). Nine (29%) the 31 azithromycin-decreased microbially-produced odd-chain lipid compounds [4 phospholipids, 2 fatty acids, 2 sterol esters, and 1 sterol lipid] also were decreased by tylosin in the mice (**Figure 3E**,**4D**). Thirteen (65%) of the 20 azithromycin-decreased microbially-produced even-chain lipid compounds [7 phospholipids, 3 fatty acids, and 3 sterol esters] also were decreased by tylosin in the mice (**Figures 3E**,**4D**). A notable difference between observations from NOD mouse cecal contents and human fecal experiments is that 10 of 11 microbially-produced sulfonolipids (SL) (**Figures 2E**) were significantly increased by tylosin in NOD mice (**Figure 3E**), while all 11 microbially-produced sulfonolipids were significantly decreased by azithromycin in humans (**Figure4D**). Thus, microbial odd-chain sulfonolipid production was highly affected by the antibiotic treatment and the initial microbial population in their hosts.

### Confirmation of 4 phospholipid compounds in NOD mouse cecal contents and in specific gut microbiota

Since phospholipids were the major antibiotic-perturbed GMPLs in both NOD mice and humans, particularly the lysophosphatidylglycerols (LPG) and phosphatidylglycerols (PG) (**Figures 3E**,**4D**), we investigated whether specific compounds could affect host responses related to T1D onset. Based on their defined chemical structures and suitability for *in vitro* and *in vivo* experiments, we selected four identified phosphatidylglycerol candidates, including three LPGs odd-chain [LPG(13:0)], and even-chain [LPG(16:0) & LPG(18:0)] and one double-chain PG [PG(15:0_15:0)] **(Table S3)** for further evaluation. The identified structure of each in mouse cecal contents was identical to the compound standard by MS/MS analysis (**Figures 5A**,**S6A; 5F**,**S6B, 5K**,**S6C, 5O**,**S6D**). LPG(13:0) showed significantly higher abundance in the cecal contents of conventional mice than in GF mice and in diet identifying it as a GMPL without abundance changes with 1P or CMT in NOD mice, but significantly decreased by azithromycin in human subjects (**Figure5B**). Azithromycin significantly decreased the abundance of PG(15:0_15:0) in the human volunteers (**Figure 5G**). For LPG(16:0) and LPG(18:0), their abundances in mouse cecal contents and in human feces were significantly decreased by the macrolide antibiotics (**Figure 5LP**). LPG(16:0) was detected in all six representative bacterial strains we had earlier tested (**Figure S2**), while LPG(18:0) were present in 5 of 6 (**Figure 5T**), but LPG(13:0) and PG(15:0_15:0) were absent. These findings indicate that the effects of antibiotic exposure on gut lipid profiles could be complex and strain-specific.

**Figure 5.**
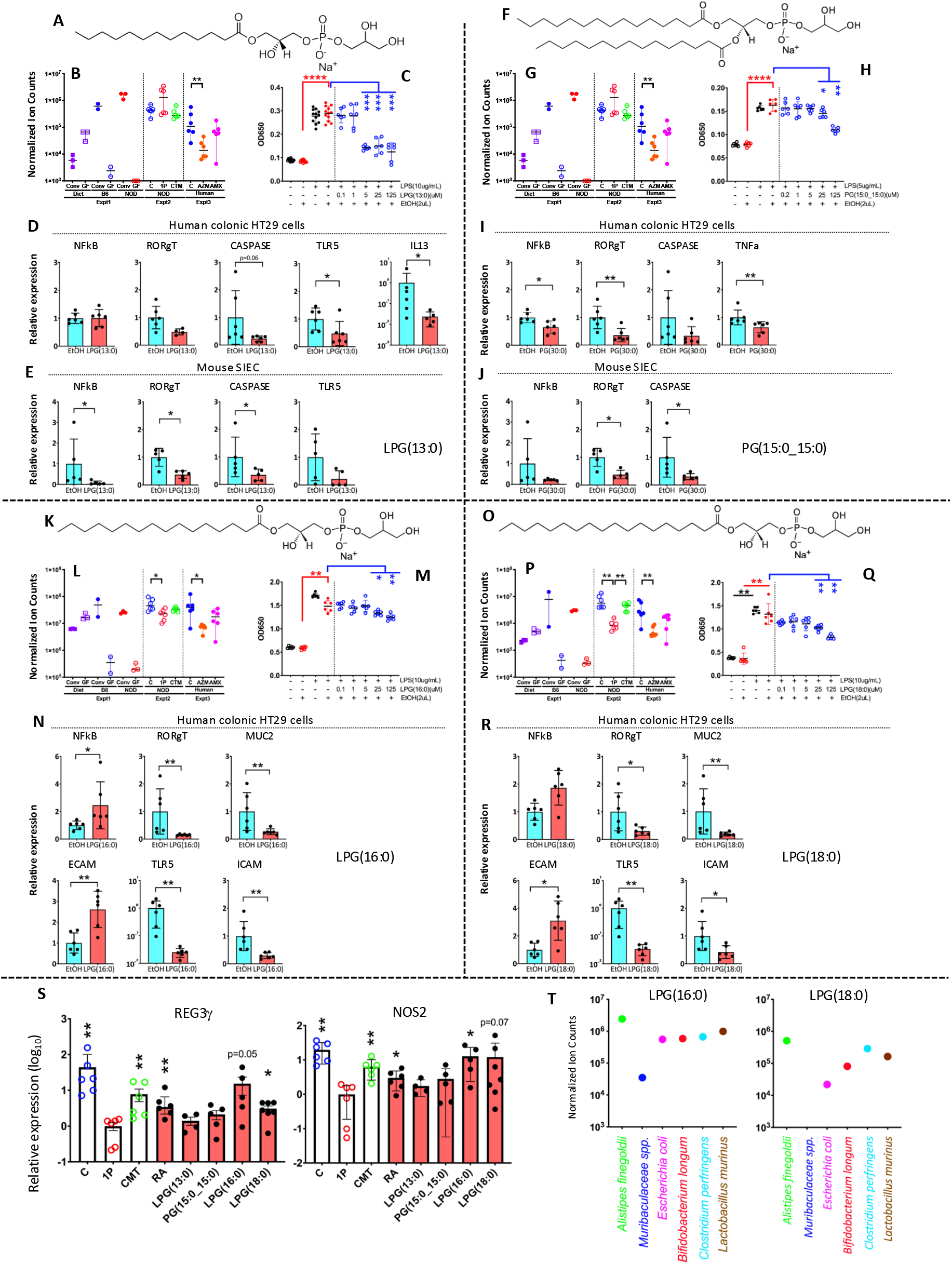
Effects of four microbially-produced phospholipids on host immune responses *in vitro* and *in vivo*. Panels A, F, K, O: Chemical structure of LPG(13:0), PG(15:0_15:0), LPG(16:0), and LPG(18:0), respectively. **Panels B, G, L, P:** Normalized abundances of the four compounds, respectively, in the three experiments (see **Figure 2**). *p<0.05; **p<0.01; Mann-Whitney *U*-test. **Panels C, H, M, Q:** Suppressive effects of the four compounds, respectively, on LPS-induced NF*κ*B activity in mouse macrophage Raw Blue cells, respectively. *p<0.05; **p<0.01; ***p<0.001; ****p<0.0001; Mann-Whitney *U*-test. **Panels D, I, N, R:** Effects of the four compounds, respectively, on expression of particular innate immune genes in human colonic epithelial HT-29 cells during co-culturing, respectively. *p<0.05; **p<0.01; Mann-Whitney *U*-test. **Panels E, J:** Effects of LPG(13:0) and PG(15:0_15:0) on the expression of particular innate immune genes in primary small intestinal epithelial cells from male NOD mice at P23. *p<0.05; **p<0.01; Mann-Whitney *U*-test. **S) In vivo effects:** RT-qPCR-based ileal gene expression of T1D early markers (REG3*γ*, NOS2) of NOD mice exposed to early-life antibiotics and then administered defined lipid compounds for restoration. Mean±SD of log_10_ values of relative expression shown. *p<0.05 compared with 1P group; Mann-Whitney *U*-test. **T)** Verification of LPG16:0 **(Left)** & LPG18:0 **(Right)** by MS2 with standards in cell pellets from six bacterial species belonging to four major mammalian intestinal phyla. *Alistipes finegoldii* strain 23-018 (Bacteroidota), *Muribaculaceae spp*. strain MH8C (Bacteroidota), *Bifidobacterium longum* strain HMXZ001 (Actinomycetota), *Escherichia coli* strain Nissle 1917 (Pseudomonadota), *Clostridium perfringens* strain S107 (Bacillota), and *Lactobacillus murinus* strain 21-100 (Bacillota).

### The phosphatidylglycerol compounds affect innate responses *in vitro*

Our prior results showed that early-life antibiotic exposure accelerated T1D onset by dysregulating genes involved in host intestinal innate immune responses^15^. Here, we investigated whether any of the 4 confirmed phospholipids affected transcription of NF*κ*B, the master regulator of innate immune responses^18^. Incubating the phospholipids alone with mouse Raw Blue macrophage NF*κ*B-activity-reporter cell lines, did not alter NF*κ*B activity (**Figure S6JKLM**). However, in the presence of *E. coli* LPS, each phospholipid significantly suppressed the LPS-induced NF*κ*B activity in a dose-dependent manner (**Figure 5CHMQ**), consistent with anti-inflammatory effects through NF*κ*B pathways, paralleling the effects of retinoic acid, a known anti-inflammatory lipid^19^ (**Figure S6H**).

Next, we evaluated whether these compounds could affect innate responses in gut epithelial cells, including human colonic epithelial HT29 cells and also with isolated mouse small intestinal epithelial cells (SIEC). Each of the compounds had a different pattern of suppressing genes related to innate immune effectors, and in some cases up-regulating them (**Figure 5DEIJNR**); these differences reflect the complex interactions of the lumen-facing epithelia of the intestine.

### Oral administration of phosphatidylglycerol compounds restores markers of intestinal immune function dysregulated by antibiotic exposure

Finally, we performed experiments to assess whether directly administering the phosphatidylglycerol compounds to mice could counteract the effects of the antibiotic-induced dysbiosis (**Figure 1D, Bottom Panel**). NOD mice that had received early-life 1P antibiotic-treatment were fed a single compound to assess whether they could reverse the 1P-induced alterations in early T1D marker ileal genes. Compared to control pups without antibiotic exposure, the 1P treatment decreased ileal expression of REG3*γ* and NOS2, two early biomarkers of T1D onset (**Figure 5S**), consistent with our prior findings^14,15^. CMT from healthy mouse donors, restored the decreased expression of REG3*γ* and NOS2 induced by 1P in the ileum (**Figure 5S**). Both LPG(16:0) and LPG(18:0) increased REG3*γ* and NOS2 expression in the ileum. Consistent with the *in vitro* data, these studies provide preliminary *in vivo* evidence of the bioactivity of these microbial compounds in the intestinal milieu and their potential to restore perturbed signaling after an antibiotic course.

### Control of bacterial production of phosphotidylglycerols

Bacterial biosynthesis of PG typically involves a two-step process catalyzed by phosphatidylglycerophosphate synthase (PgsA) and phosphatidylglycerophosphate phosphatases (PgpA, PgpB, PgpC) (**Figure S7A**), while lysophosphatidylglycerol (LPG) is typically formed by the partial hydrolysis of PG, catalyzed by phospholipase A2 (PLA2)^26,28^. A novel alternative pathway for PG synthesis in *E. coli* is PgsA-independent, involving the conversion of phosphatidylethanolamine and glycerol into PG, catalyzed by ClsB, a phospholipase D-type cardiolipin synthase^29^. The β-ketoacyl-ACP synthase III (KASIII) enzyme (FabH) plays a key role in determining fatty acid structure in bacteria (**Figure S7A**), including whether the fatty acid side chain of lipids like LPG/PG is odd-chain or even-chain^17^. That the relative abundances of these genes in the fecal microbiome was significantly affected by an early-life single dose of macrolide antibiotic (tylosin) and restored by CMT in NOD mice, and that the abundances of these genes could be substantially controlled by administrating azithromycin in humans (**Figure 4F, S7B, C**), provides opportunities to regulate the pro- and anti-inflammatory milieu in the intestine for therapeutic purposes.

## Discussion

Our prior studies of NOD mice showed that antibiotic exposure induced gut microbiome perturbation in early life which accelerated T1D onset, and that this could be restored by maternal cecal microbiota transplant (CMT) through regulating host innate immune responses^13-15^. Based on clues from metagenomic analyses of cecal microbiome which highlighted differentiation of bacterial lipid metabolism pathways during microbiota perturbation, here we systematically characterized gut microbiota lipid profiles in conventional and germ-free mice and now identified a group of specific microbiota-produced lipid compounds. Using the pipeline that comparing lipidomics of gut microbiota from antibiotic-treated mice and humans and their untreated controls as a strategy for discovery, from >700 identified lipids, we identified a group of gut bacterial-derived lipids that had been reduced by the antibiotic treatments, and sometimes restored by the CMT. We then focused on a subset of these defined lipid compounds with defined structures to further investigate their biological activities *in vitro* and *in vivo*. Further work is warranted to define optimal concentrations and combinations, and to examine efficacy in animal models of gut inflammatory disorders such as T1D, or Inflammatory Bowel Disease (IBD) for preventive or therapeutic uses.

Bacterial-derived lipids also include highly diverse and abundant phospholipids such as LPG, PG, LPC, PC, and CL, which were released from bacterial cell membranes or outer membrane vesicles (OMVs) and could be critical in mediating host immunity^20-24^. The structural diversity of phospholipids not only reflects the biochemical and biophysical characteristics of bacterial membranes enabling bacterial adaptation in the complex gut environment, but also could differentially mediate host-microbe communication and modify host immune responses^20,21,25,26^. We focused on four defined phospholipids, LPG(13:0), LPG(16:0), LPG(18:0), and PG(15:0_15:0) whose abundance consistently varied between microbiota samples from antibiotic-treated and control subjects, representing the potential of the broader group for host interactions. Our results suggest that these bacterial lipids could serve as therapeutic agents for the prevention or treatment of inflammatory and immune-mediated processes originating in the gut, including T1D and IBD^27^.

## Methods

### Lipid compound standards

SPLASH LipidoMIX Mass Spec Standard, including PC(15:0_18:1-d7), PE(15:0_18:1-d7), PS(15:0_18:1-d7), PG(15:0_18:1-d7), PI(15:0_18:1-d7), PA(15:0_18:1-d7), LPC(18:1-d7), LPE(18:1-d7), cholesteryl ester(18:1-d7), MG(18:1-d7), DG(15:0_18:1-d7), TG(15:0_18:1-d7_15:0), SM(18:1-d7), and cholesterol(d7)] were purchased from Avanti Polar Lipids (cat# 330707, Birmingham, AL) and used as internal standards for lipidomic MS analysis. For use as references for specific cecal lipid MS2 analyses and for both in vitro and in vivo experiments, we purchased Lysophosphatidylglycerols LPG (13:0) [1-tridecanoyl-sn-glycero-3-phospho-(1’-rac-glycerol) (sodium salt)], LPG (16:0) [1-palmitoyl-2-hydroxy-sn-glycero-3-phospho-(1’-rac-glycerol) (sodium salt)], & LPG (18:0) [1-stearoyl-2-hydroxy-sn-glycero-3-phospho-(1’-rac-glycerol) (sodium salt)], and phosphatidylglycerol PG (15:0_15:0) [1,2-dipentadecanoyl-sn-glycero-3-phospho-(1’-rac-glycerol) (sodium salt)] (Avanti Polar Lipids), and retinoic acid (RA) and *E. coli* lipopolysaccharides (LPS) was purchased from Sigma-Aldrich (cat# L2012-10MG, Saint Louis MO).

### Preparation of lipids

Total lipids were extracted using the methyl tert-butyl ether (MTBE) method . Fresh extraction solvent was prepared on the same day of total lipid extraction by mixing 0.1 M hydrochloric acid (Sigma-Aldrich cat# 1090601000), methanol (Sigma-Aldrich cat# 34860), and the SPLASH LipidoMIX Internal Standard in a 500:495:5 v/v/v ratio. For each 10-15 mg sample of cecal contents, feces, or mouse diet, 50 µL fresh extraction solvent was added to 2 mL screw cap tubes containing 200 µL of 0.1 mm glass beads (BioSpec cat# 11079101, Bartlesville OK). The samples were homogenized at 2000 rpm for 45 seconds, then paused for 45 seconds, repeating for 5 cycles to avoid overheating. The mixture was then combined with 2 volumes of MTBE, vortexed for 30 seconds, and left to stand at room temperature for 1 minute to allow layer separation. The top MTBE layer containing total lipids (600 µL) from each sample was collected and air-dried in a fume hood for ∼16 hours. The dried samples were stored at - 80°C until mass spectrometry analysis performed at the Metabolomics Shared Resource, Rutgers Cancer Institute. The dried total lipid samples were resuspended in 150 µL of resuspension solvent (isopropanol mixed with methanol at 1:1 v/v), followed by centrifugation at 13,000 g for 10 minutes. The top layer (100 µL) was then transferred to glass autosampler tubes.

### Liquid chromatography–mass spectrometry analysis

Reverse phase separation was performed on a Vanquish Horizon UHPLC system (Thermo Fisher Scientific, Waltham MA) with a Poroshell 120 EC-C18 column (150 mm × 2.1 mm, 2.7 μm particle size, Agilent InfinityLab, Santa Clara CA) using a gradient of solvent A (90%:10% water : methanol with 34.2 mM acetic acid, 1 mM ammonium acetate), and solvent B (75%:25% isopropanol : methanol with 34.2 mM acetic acid, 1 mM ammonium acetate). The gradient was 0 min, 25% B; 2 min, 25% B; 5.5 min, 65% B; 12.5 min, 100% B; 19.5 min, 100% B; 20.0 min, 25% B; 30 min, 25% B. The flow rate was 200 μl/min. The injection volume was 5 μL and the column temperature was 55 °C. The autosampler temperature was set to 4°C and the injection volume was 5µL. The full scan mass spectrometry analysis was performed on a Thermo Q Exactive PLUS with a HESI source which was set to a spray voltage of -2.7kV under negative mode and 3.5kV under positive mode. The sheath, auxiliary, and sweep gas flow rates were 40, 10, and 2 (arbitrary unit) respectively. The capillary temperature was set to 300°C and the aux gas heater was 360°C. The S-lens RF level was 45. The m/z scan range was set to 100 to 1200 m/z under both positive and negative ionization modes. The AGC target was set to 1e6 and the maximum IT was 200 ms. The resolution was set to 140,000 at m/z 200. All samples were analyzed by positive and negative electrospray ionization (ESI+/ESI-) in full scan MS mode. Quality control samples (pooled QC) were prepared by combining equivalent volumes of each sample and were used in data-dependent MS-MS (ddMS2) acquisition for lipid identification purposes.

Targeted lipidomics for confirming Iipid identity were performed by Parallel Reaction Monitoring (PRM) acquisition mode. The PRM parameters were as follows: default charge, 1; resolution 17,500 at m/z 200; isolation window, 2 m/z; AGC target was set to 1e5 and the maximum IT was 100 ms. Stepped normalized collision energy (NCE) was set to 20, 30, and 40.

### Lipidomic Data Processing

Thermo RAW data files were acquired using Xcalibur 4.3 software (Thermo Scientific) and were converted to ABF format using the ABF converter (accessible at: http://www.reifycs.com/AbfConverter). The lipid annotation was performed using MS-DIAL 4.9 (http://prime.psc.riken.jp/compms/msdial/main.html) and the lipid quantitation was performed using El-MAVEN. Normalization was done based on the Splash® Lipidomix® internal standards.

### Mice

The NOD/ShiLtJ Germ-free mice were obtained from Kathy D. McCoy laboratory of University of Calgary and C57BL/6J Germ-free mice were purchased from Charles River Laboratories (Wilmington DE), and were bred and maintained in a gnotobiotic facility at Rutgers New Jersey Medical School with sterilized chow diets (LabDiet 5LG4, LabDiet, St. Louis MO). Conventional NOD/ShiLtJ mice and C57BL/6J mice were purchased from Jackson Laboratory (Bar Harbor ME) and bred in an SPF vivarium at Rutgers University’s School of Public Health animal facility and at the Robert Wood Johnson Medical School Research Tower animal facility, with chow diets (PicoLab 5058, LabDiet). Both germ-free and conventional mice (NOD/ShiLtJ and C57BL/6J) in the above facilities were maintained in the same environment of 23 °C ± 1 °C, on a 12:12-hour light-dark cycle. All animal procedures were approved by the Rutgers University Institutional Animal Care and Use Committee (IACUC protocols no. 201900013, 201900017 and 201900032).

### Human fecal specimens

Fecal specimens were obtained from young adults (age 23-39 years) enrolled in the Microbial, Immune, and Metabolic Perturbations by Antibiotics (MIME) Study. Subjects received amoxicillin (AMX, 500 mg every 12 hours for 7 days), or azithromycin (AZM, 500 mg on day 1, followed by 250 mg every 24 hours for 4 days), or no antibiotic (Control). Each treatment group had 6 subjects; 3 males/3 females. The fecal samples were collected and directly frozen at -20°C at participants’ homes, and then transported in dry ice to study laboratories at NIH, and subsequently to Rutgers University, where they were frozen at -80°C for microbiome and lipidomic analysis. All procedures were approved by the NIH Institutional Review Board (IRB protocol 16-I-0078).

### Microbe strains

All bacterial strains were grown at 37°C in an anaerobic Chamber (Coy Lab Products, Grass Lake MI) under an atmosphere of 90% N2, 5% CO2, and 5% H2 for 24-72 h. *Alistipes finegoldii* strain 23-018 (isolated from P23 NOD mouse cecal contents in Blaser Lab) was cultured in Brain Heart Infusion (BHI) medium (Thermo Scientific, Waltham MA), *Muribaculaceae spp*. strain MH8C (gift from the Carolina Tropini Lab) and was cultured on Tryptic soy agar (TSA) with 5% sheep blood agar medium (Thermo Scientific), *Escherichia coli* strain Nissle 1917 was cultured in Luria-Bertani (LB) broth medium, *Bifidobacterium longum* strain HMXZ001 (isolated from ECAM baby feces in Blaser lab) was cultured in Bifidobacterium Selective Medium (BSM) (Millipore Sigma, Burlington MA), *Clostridium perfringens* strain S107 (ATCC, Manassas VA) were cultured in Reinforced Clostridial Medium (RCM) (Thermo Scientific), and *Lactobacillus murinus* strain 21-100 (isolated from P23 NOD mouse cecal contents in Blaser Lab) were cultured in De Man, Rogosa, and Sharpe (MRS) medium (Thermo Scientific).

### Sample 16S Microbiome analysis and Statistical methods

Fecal microbiota DNA from 18 subjects was extracted using the DNeasy PowerSoil-htp HTP 96 Kit (Qiagen, Hilden, Germany). The V4 region of bacterial 16S rRNA genes was amplified in triplicate reactions using barcoded fusion primers 515F/806R, which amplifies bacterial and archaeal 16S genes. The DNA concentration of the V4 amplicons for each sample was measured using the Quant-iT PicoGreen dsDNA assay kit (Life Technologies, Eugene OR). Samples were pooled in equal quantities. These set pools were then purified using the Qiaquick PCR purification kit (Qiagen) to remove primers, quantified using the high-sensitivity dsDNA assay kit and the Qubit 2.0 Fluorometer (Life Technologies) and then combined at equal concentrations to form the sequencing library. About 254 bp V4 region was sequenced using the Ilumina MiSeq 2 × 150 bp platform. Quantitative insights for microbial ecology (QIIME, Version Qiime 2-2022.8) were used for quality filtering and downstream analysis for α-diversity (Shannon index), β-diversity (Jaccard distance matrix) and compositional analysis. Sequences were filtered for quality trimmed, de-noised, merged and then the chimeric sequences were removed using DADA2 plugin to generate the feature table. Taxonomy was assigned using Silva 138. Significant differences in alpha diversity between experimental groups were determined using Kruskal-Wallis method, while differences in β-diversity were tested by pair-wise PERMANOVA with 999 permutations.

### Cell culture and co-culture assays

Murine macrophage NF*κ*B-SEAP reporter RAW-Blue cells (InvivoGen Cells, cat# raw-sp, San Diego CA) were grown in Dulbecco Modified Eagle Medium (DMEM) (Corning, Tewksbury MA) supplemented with 4.5% glucose, 10% fetal calf serum (FCS) (Corning), 2 mM L-glutamine, 1 x Penicillin-streptomycin (Gibco, Waltham MA), and 100 µg/mL zeocin (InvivoGen Cells) in a humidified incubator with 5% CO2 at 37°C. The cells were passaged when they reached 70% confluence for 6-10 generations. Cells were scraped, resuspended in fresh media, and plated in a flat-bottom 96-well tissue culture plate (Corning) at final density 2 x 10^5^ cells/well in 180 µL. The cells were incubated for 2 hours at 37°C in an atmosphere of 5% CO2, then treated with the target lipid compound at a final concentration from 0.1-125 µM in 1% (v/v) EtOH or with a blank reagent (EtOH) with final concentration 1% (v/v) as a control. After 1 hour of incubation, the lipid-cell co-cultures were treated with *E. coli* LPS (Sigma -Aldrich) to a final concentration of 10 µg/mL. After 24 hours of stimulation, supernatants were collected, and NF*κ*B activation determined using the detection medium QUANTI-Blue, prepared according to the manufacturer’s recommendations.

Human colonic epithelial cell line HT-29 (ATCC, Manassas VA) was cultured in RPMI 1640 medium (Corning) with 10% FCS and 1x penicillin-streptomycin in a humidified incubator with 5% CO2 at 37°C and passaged when they reached 90% confluence. For co-culturing with lipid compounds, the cells were scraped and resuspended in fresh media, and plated into 6-well tissue culture plates (Corning) and grown to >90% confluence in RPMI 1640 medium with 10% FBS without antibiotics (2 mL per well) and incubated for 1 hour to improve attachment, then each lipid compound added at a final concentration of 50 µM in 1% EtOH, or blank reagent EtOH as a reference control, incubating for 16 hours at 37°C in 5% CO2. The attached cells were washed with ice-cold PBS, resuspended in TRIzol reagent (Qiagen), and frozen at -80°C for RNA extraction.

To isolate mouse small intestinal epithelial cells (SIEC), male NOD mice were sacrificed at P23, the small intestine was removed and opened longitudinally, rinsed with cold HBSS (Gibco) to remove contents, and cut into 2-cm pieces, and incubated on ice in 20 mL DMEM (Corning) supplemented with 10% FCS (Gibco) with Collagenase type I (Sigma-Aldrich) 200 U/mL at 37°C for 30 min, with gently shaking at 100 rpm. The epithelial layer was gently dissociated by pipetting up and down from intestinal tissue and collected, filtered through a 100 µm cell strainer (Fisher Sci, Waltham MA) and centrifuged at 300 g for 5 min to pellet cells. Cells pooled from 3 mice were cultured and passaged in DMEM medium (Corning) supplemented with 10% FCS and 1x penicillin-streptomycin. For incubation with lipid compounds, fresh sub-cultured cells were seeded at high density into 12-well tissue culture plates (Corning) in DMEM medium with 10% FBS without antibiotics (1 mL per well), cultured to >90% confluence for 1 hour, then incubated in a 2 mL co-culture system for 16 hours at 37°C in 5% CO2 with each lipid compound (4 µL) at a final concentration of 50 µM in EtOH, or blank reagent EtOH as a reference control. The attached SIEC cells were washed with ice-cold PBS, resuspended in TRIzol and frozen at -80°C for RNA extraction.

### RT-qPCR for host target gene expression

Total RNA was extracted from TRIzol-collected cells using the QIAgen Mini RNeasy kit (Qiagen) and cDNA was synthesized from the total RNA samples using the Verso cDNA kit (Thermo Scientific) according to the manufacturers’ instructions. qPCR was performed on a LightCycler 480 system (Roche, Branchburg NJ) using 10 ng of synthesized cDNA, target gene-specific primer pairs (**Table S4**), and Power SYBR Green PCR Master mix (Roche). Target mRNA levels were normalized to 18S rRNA or the housekeeping gene GAPDH as internal controls for each sample. For group mean comparisons, the Mann-Whitney t-test was performed, with p-value < 0.05 indicating significance.

### Oral administration of lipids to mice

To evaluate the *in vivo* effects of administering specific lipid compounds to young NOD mice, dams and their litters were randomly assigned to control (C) or 1PAT antibiotic (1P) groups as previously described^15^. A therapeutic dose of the macrolide tylosin tartrate (Sigma-Aldrich, Billerica MA) was given to 1P pups in their non-acidified drinking water at 333 mg/L on P5-P10. At postnatal day (P) 12, pups from 3 litters in the 1P group received a single gavage of fresh prepared lipid compounds LPG(13:0), PG(15:0_15:0), LPG(16:0), or LPG(18:0), respectively, at a dose of 0.3 mg/kg body weight (60 µL) daily for one week to P18. As controls, pups in the C group received the same volume of the blank solvent reagent (PBS-2% EtOH). Additional litters of 1P mice received: (i) cecal material transfer (CMT) from a pool of cecal contents from 3 male NOD mice at P23; or (ii) retinoic acid (3 mg/kg body weight) in a single daily dose from P12 to P18. From each litter, four pups (2 male/2 female) were sacrificed on P19, and ileum and colon collected in RNAlater for RT-qPCR-based evaluation of expression of host genes associated with T1D development. Cecal contents were also collected and frozen for lipidomic analysis and 16S rRNA gene sequencing.

## ACKNOWLEDGEMENTS

These studies were supported by U01 AI 22285 from the National Institutes of Health, and from the TransAtlantic Program of the Fondation Leducq (33.17CVD01), and by the Zlinkoff and Emch Funds. The Rutgers Cancer Institute Metabolomics Shared Resource is supported, in part, with funding from NCI-CCSG P30CA072720-6852. We thank Kathy D. McCoy (U Calgary), Carolina Tropini (U British Columbia), Gang Fang (Mount Sinai Medical Center) for mice and bacterial strains, and the Rutgers Gnotobiotic core facility for assistance with these studies.

