## Supplemental materials for "Gut microbiota phospholipids regulate intestinal gene expression and can counteract the effects of antibiotic treatment"

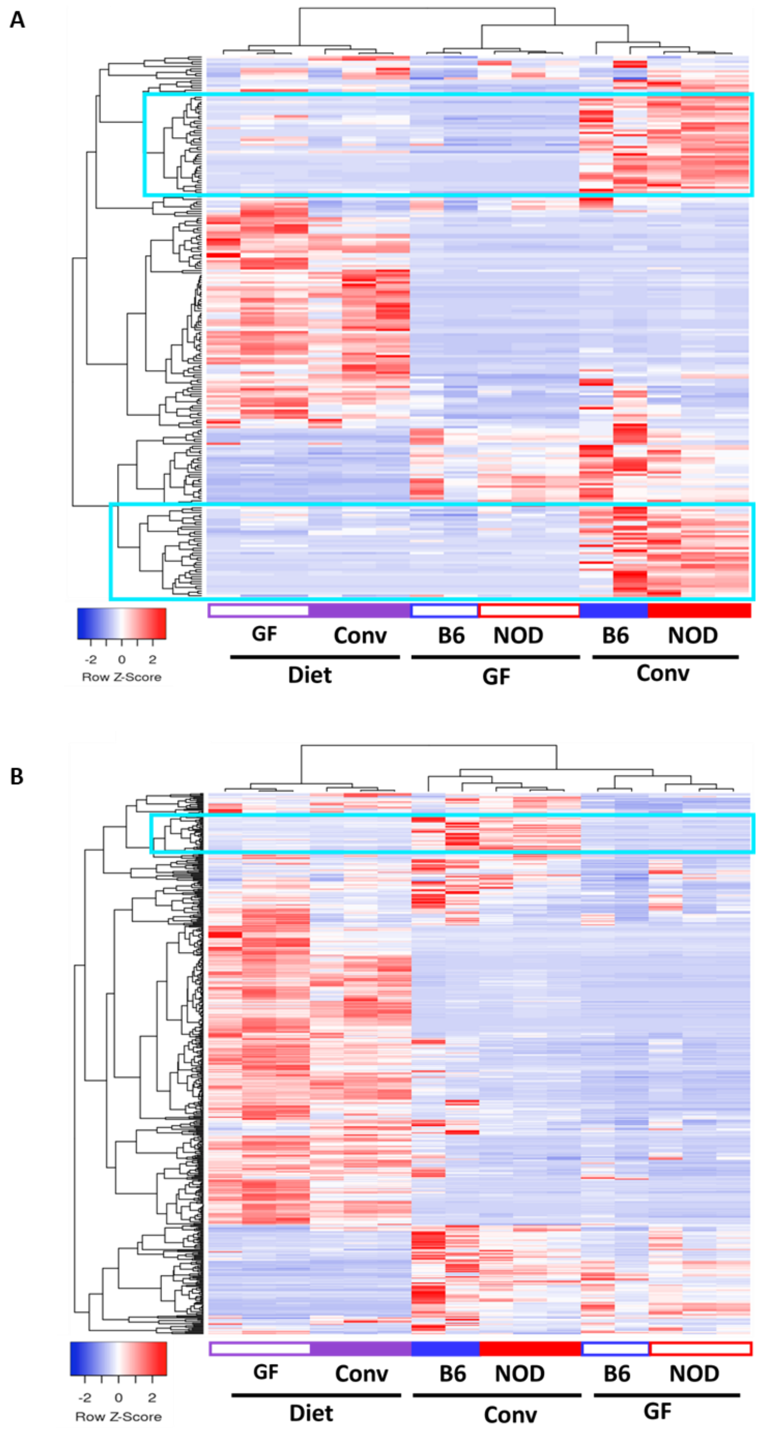


**Supplemental Figure 1. Clustering of lipid compounds in diet and conventional and germ-free mice in Experiment 1.** Unsupervised hierarchical clustering based on 230 odd-chain lipids **(A)** and 517 even-chain lipids **(B):** Blue boxes indicate microbially-produced lipid compounds, highly abundant in cecal contents from conventional mice but low in diets and low in cecal contents from germ-free mice.

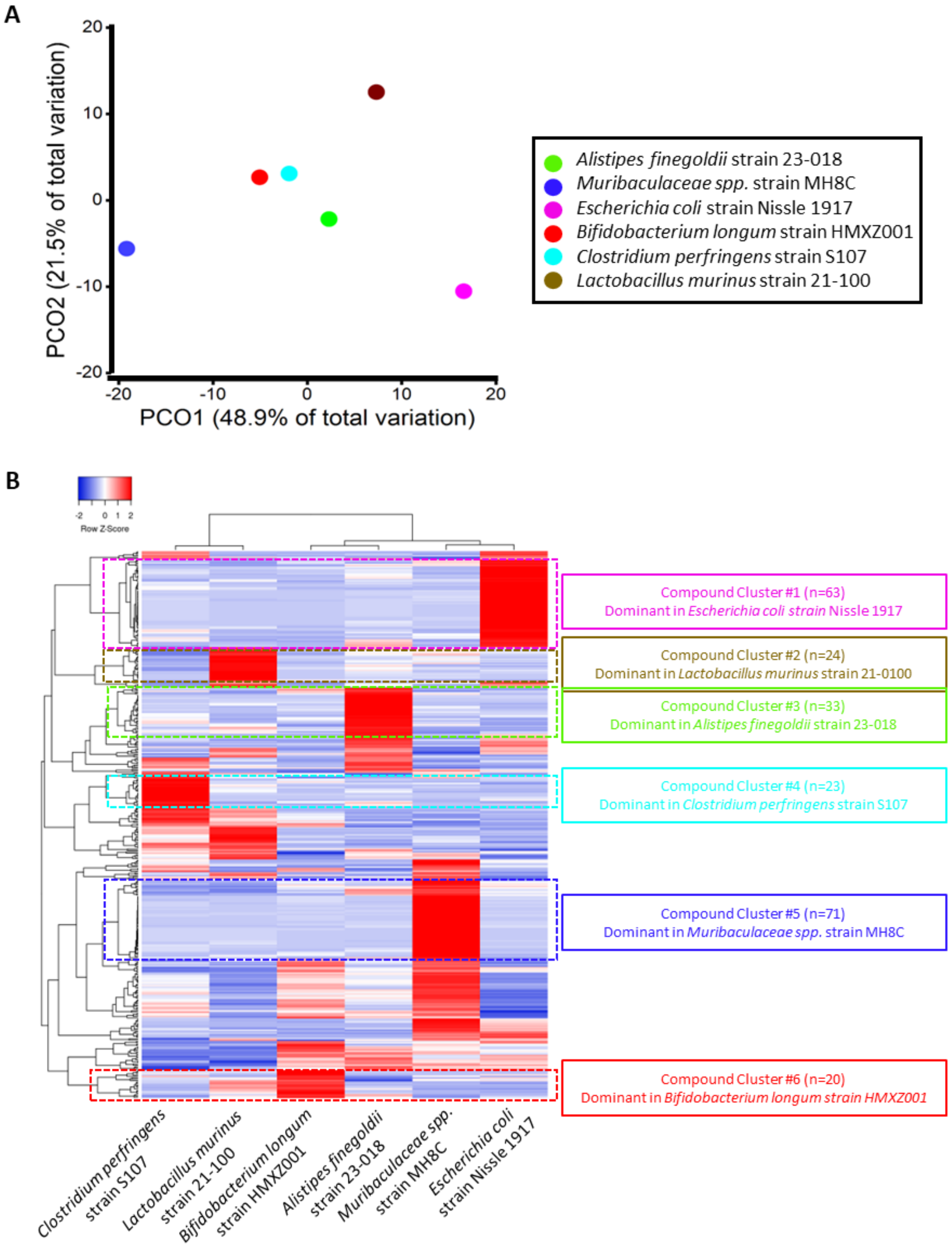


**Supplemental Figure 2. Non-redundant lipid compounds in 6 representative intestinal bacterial species.** Verification of defined compounds by MS2 with standards in cell pellets from six bacterial species belonging to the four major phyla in the intestine. **A)** Bray-Curtis distance based on the 391 non-redundant lipid compounds identified in the cell pellets of the 6 bacterial species. **B)** Unsupervised hierarchical clustering based on 391 nonredundant lipid compounds across the six strains. Strain-specific lipid compound clusters are highlighted within the boxes.

### 1P-significantly altered microbially-produced lipids identified in volcano plot (n=36)

| Compound | $Log_2$(FD) (1P/C) | p.Value ($-log_{10}$) |
|---|---|---|
| FA(19:0;O) | -3.6 | 3.19 |
| FA(20:0;O) | -3.8 | 5.32 |
| FA(21:0;(2OH)) | -3.2 | 4.43 |
| **FA(22:0;O)** | **-4.0** | **5.16** |
| **FA(22:1;(2OH))** | **-4.1** | **4.72** |
| **FA(22:3)** | **-5.1** | **4.28** |
| FA(23:0;O) | -3.3 | 4.44 |
| **FA(23:1;O)** | **-3.9** | **4.89** |
| FA(24:0;(2OH)) | -2.4 | 4.33 |
| **FA(24:1;O)** | **-4.1** | **5.08** |
| FA(24:2;O) | -4.5 | 5.82 |
| FA(25:1;O) | -2.6 | 4.29 |
| FA(32:9) | -3.6 | 4.61 |
| HexCer(18:1;2O/16:0) | -3.1 | 2.95 |
| **LPG(15:1)** | **-2.3** | **2.71** |
| **LPG(18:0)** | **-2.9** | **4.64** |
| **LPG(O-18:2)** | **-3.0** | **3.15** |
| MG(21:4) | -7.9 | 9.43 |
| **MG(21:5)** | **-5.5** | **7.01** |
| **PG(16:0_18:0)** | **-2.0** | **3.48** |
| **PG(18:0_18:1)** | **-2.7** | **4.77** |
| **PG(O-18:2_16:0)** | **-3.5** | **4.78** |
| **PI(15:0_15:0)** | **-3.4** | **2.60** |
| **PS(31:0)** | **-3.5** | **2.63** |
| **SE(24:1;O4/17:0)** | **-2.2** | **4.47** |
| **SE(24:1;O4/18:0)** | **-2.6** | **3.26** |
| **SE(24:1;O4/18:1)** | **-2.7** | **2.65** |
| **ST(27:1;O)** | **-3.3** | **4.61** |
| ST(29:1;O) | -3.9 | 5.89 |
| TG(14:0_16:0_18:0) | -3.0 | 6.82 |
| PE(15:0_18:3) | 2.1 | 3.78 |
| **SL(17:0;O/15:0)** | **2.8** | **5.46** |
| **SL(17:0;O/17:0)** | **2.1** | **5.50** |
| **SL(18:0;O/17:0)** | **3.6** | **8.72** |
| **SL(18:0;O/17:1;O)** | **3.6** | **5.72** |
| **SL(19:0;O/17:1;O)** | **4.6** | **6.32** |

### 1P-significantly altered host-produced lipids (n=3)

| Compound | $Log_2$(FD) (1P/C) | p.Value ($-log_{10}$) |
|---|---|---|
| ST(27:2;O) | -2.1 | 4.60 |
| DG(37:7) | 2.6 | 3.17 |
| SL(17:0;O/17:1;O) | 3.0 | 3.23 |

### 1P-significantly altered diet-enriched lipids (n=156)

| Compound | $Log_2$(FD) (1P/C) | p.Value ($-log_{10}$) |
|---|---|---|
| 1P-decreased (n=155) (See Table S4) | | |
| Cer(18:0;2O/12:0) | 2.6 | 4.61 |

Identified by volcano plot analysis with $log_2$FC (1P/C) <-2 or >2, and $log_{10}$-transformed p<0.01.

**Bold:** Also listed in **Figure 4F** as 1P-decreased or 1P-increased microbially-produced lipids.

**Supplemental Figure 3. Lists of significant antibiotic-exposure-altered lipid compounds in NOD mice, identified in volcano plot analysis.** In volcano plot analysis using the criteria of a geometric mean fold-difference >4 and log-transformed p-value <0.01, 36 microbially-produced compounds were significantly altered by 1P exposure, including 30 decreased and 6 increased (Panel Left); 3 host-produced compounds were altered by 1P, including 1 decreased and 2 increased (Panel Right top); and 156 diet-enriched compounds were significantly altered by 1P, including 155 decreased (See **Supplemental Table 1**) and 1 increased (Panel Right bottom).

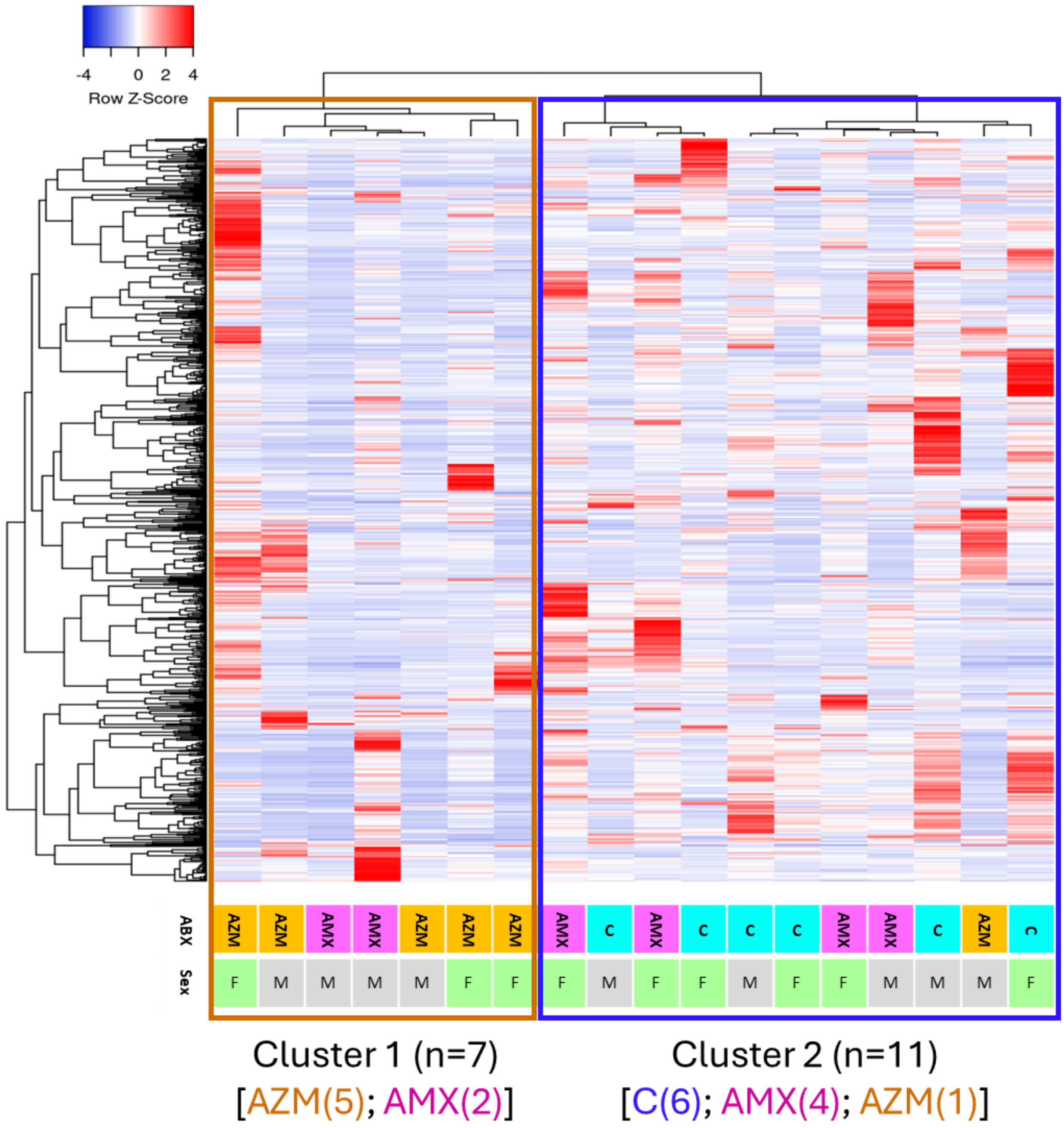


**Supplemental Figure 4. Unsupervised clustering analysis based on 728 non-redundant lipid compounds identified in fecal samples from 18 human subjects.** Samples are from three experimental groups of human volunteers: control, AZM-, and AMX-exposure. Two distinct clusters were identified among the subjects. Cluster 1 includes 5 of the 6 AZM-exposed subjects and 1 AMX-exposed subject. Cluster 2 includes all 6 control subjects, 4 of the 6 AMX-exposed subjects and 1 AZM-exposed subjects.

### AZM-significantly decreased microbially-produced lipids identified in volcano plot (n=39)

| Compound | $Log_2$(FD) (AZM/C) | p.Value ($-log_{10}$) |
|---|---|---|
| FA(19:0) | -2.1 | 2.00 |
| FA(19:0;O) | -3.3 | 2.15 |
| FA(20:0;O) | -3.4 | 2.32 |
| FA(21:0;(2OH)) | -4.5 | 2.86 |
| **FA(22:0;O)** | **-4.0** | **2.55** |
| **FA(22:1;(2OH))** | **-3.7** | **2.44** |
| FA(23:0;O) | -4.0 | 2.47 |
| **FA(23:1;O)** | **-4.9** | **2.47** |
| FA(24:0;(2OH)) | -3.6 | 2.60 |
| **FA(24:1;O)** | **-3.8** | **2.20** |
| FA(24:2;O) | -4.1 | 2.07 |
| FA(25:0;O) | -3.8 | 2.92 |
| FA(25:1;O) | -3.6 | 2.22 |
| FA(26:0;O) | -3.9 | 2.67 |
| FA(26:1;O) | -3.1 | 2.20 |
| FA(27:0;O) | -4.5 | 2.85 |
| **LPE(17:0)** | **-2.7** | **2.59** |
| **LPG(13:0)** | **-3.0** | **2.45** |
| **LPG(14:0)** | **-2.2** | **2.46** |
| **LPG(15:1)** | **-2.9** | **2.44** |
| **LPG(16:0)** | **-2.3** | **2.51** |
| **LPG(17:0)** | **-3.4** | **2.12** |
| **LPG(17:1)** | **-4.0** | **2.23** |
| **LPG(18:0)** | **-2.5** | **2.76** |
| **LPG(18:1)** | **-2.2** | **2.00** |
| **PG(16:0_16:1)** | **-2.3** | **3.12** |
| **PG(O-18:2_16:0)** | **-2.4** | **2.69** |
| **SE(24:1;O4/17:0)** | **-2.9** | **2.05** |
| **SE(24:1;O4/18:1)** | **-3.2** | **2.12** |
| **SE(24:1;O4/18:2)** | **-2.5** | **2.14** |
| **SL(16:0;O/15:0)** | **-6.1** | **2.11** |
| **SL(17:0;O/15:1;O)** | **-6.9** | **2.38** |
| SL(32:1;O) | -8.2 | 5.37 |
| **SL(17:0;O/16:0)** | **-5.1** | **2.34** |
| **SL(17:0;O/16:1;O)** | **-5.1** | **2.33** |
| **SL(33:1;O)** | **-4.2** | **2.52** |
| **SL(17:0;O/17:0)** | **-5.4** | **2.36** |
| **SL(18:0;O/17:1;O)** | **-5.2** | **2.88** |
| **SL(19:0;O/17:1;O)** | **-4.9** | **2.22** |

No microbially-produced compound
Were increased by AZM significantly

### AZM-significantly altered host-produced lipids (n=5)

| Compound | $Log_2$(FD) (AZM/C) | p.Value ($-log_{10}$) |
|---|---|---|
| FA(16:1;3O) | -3.1 | 2.48 |
| FA(36:1) | -2.2 | 2.27 |
| SL(17:0;O/17:1;O) | -4.5 | 2.66 |
| ST(27:0;O;S) | -3.2 | 2.30 |
| HexCer(18:0;3O/20:0;(2OH)) | 2.9 | 2.44 |

### AZM-significantly altered diet-enriched lipids (n=11)

| Compound | $Log_2$(FD) (AZM/C) | p.Value ($-log_{10}$) |
|---|---|---|
| Cer(18:2;2O/26:6;O) | -3.0 | 2.51 |
| DG(20:0) | -2.2 | 2.17 |
| FA(14:0) | -2.7 | 2.47 |
| FA(18:1) | -2.5 | 4.69 |
| FA(18:2) | -2.0 | 2.03 |
| FA(18:3;O) | -2.4 | 2.35 |
| FA(19:0;2O) | -2.2 | 2.36 |
| FA(20:1) | -2.8 | 3.99 |
| NAE(19:4) | -6.5 | 3.07 |
| Cer(18:2;2O/26:6;O) | -3.0 | 2.51 |
| Cer(17:3;2O/38:6) | 4.1 | 2.10 |

Identified by volcano plot analysis
with $log_2$FC (AZM/C) <-2 or >2,
and $log_{10}$-transformed p<0.01.

**Bold:** Also listed in **Figure 5E** as
AZM-decreased microbially-produced lipids.

### AMX-significantly altered lipids (n=3)

| Compound | $Log_2$(FD) (AZM/C) | p.Value ($-log_{10}$) |
|---|---|---|
| DG(O-18:0_22:0)* | -3.57 | 3.52 |
| PC(38:3) | -3.78 | 2.30 |
| TG(18:2_18:2_9:0;2O)** | 2.72 | 2.17 |

* as one host-produced compound
**as one diet-enriched compound

**Supplemental Figure 5. Lists of significant antibiotic-exposure-altered lipid compounds in human subjects identified in volcano plot analysis.** In the analysis using the criteria of geometric mean fold-difference >4 and log-transformed p-value <0.01, a total of 39 microbially-produced compounds were significantly decreased by AZM, and none were increased (Panel Left); 5 host-produced compounds were altered by AZM, including 4 decreased and 1 increased (Panel Right top); and 11 diet-enriched compounds were significantly altered by AZM, including 10 decreased and 1 increased (Panel Right middle). For all 728 compounds, only 3 compounds were significantly altered by AMX, including 2 decreased (1 host-produced compound), and 1 increased (diet-enriched) (Panel Right bottom).

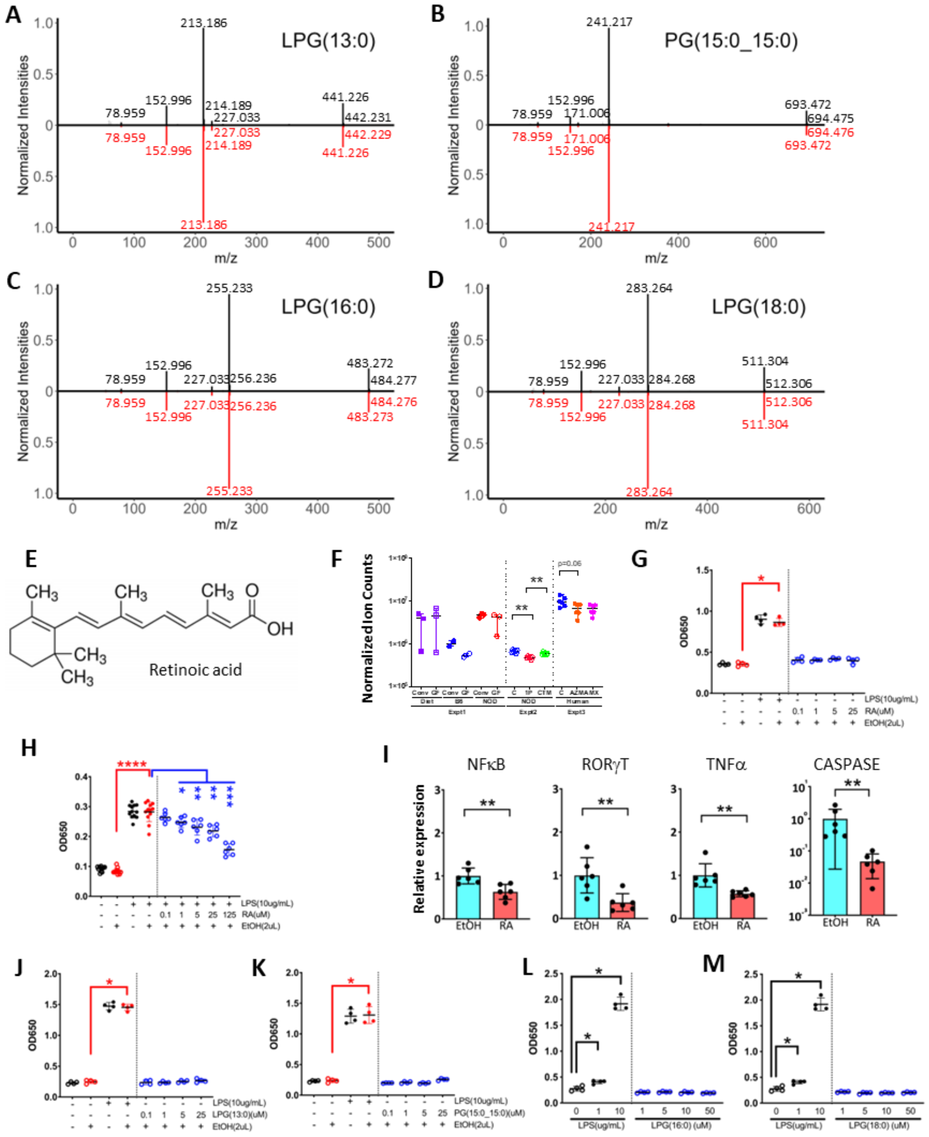


**Supplemental Figure 6. Mass spectrometric confirmation of the structure of four (lyso-)PG compounds in NOD mouse cecal contents and effects of retinoic acid and the 4 (lyso-)PG compounds on host immune responses *in vitro*. A)** LPG(13:0), **B)** PG(15:0_15:0), **C)** LPG(16:0), and **D)** LPG(18:0). The normalized intensities of detected fragments (m/z) are plotted for each lipid compound. Peaks in black above the X-axis correspond to the targeted lipid compound, while peaks in red below the X-axis represent the reference standard. **E)** Chemical structure of retinoic acid. **F)** The normalized abundances of retinoic acid in the 3 lipidomic analysis experiments. **$p<0.01$; Mann-Whitney *U*-test. **G)** No effects of retinoic acid alone on NFκB activity in mouse macrophage Raw Blue cells. **H)** Repressive effects of retinoic acid on LPS-induced NFκB activity in mouse macrophage Raw Blue cells. *$p<0.05$; **$p<0.01$; Mann-Whitney *U*-test. **I)** Significant effects of retinoic acid on the expression of particular innate immune genes in human colonic epithelial HT-29 cells during co-culture. **$p<0.01$; Mann-Whitney *U*-test. **J-M)** No effects of individual (lyso-)PG compounds on NFκB activity in mouse macrophage Raw Blue cells, in the absence of LPS co-stimulation: **J)** LPG(13:0); **K)** PG(15:0_15:0); **L)** LPG(16:0); **M)** LPG(18:0).

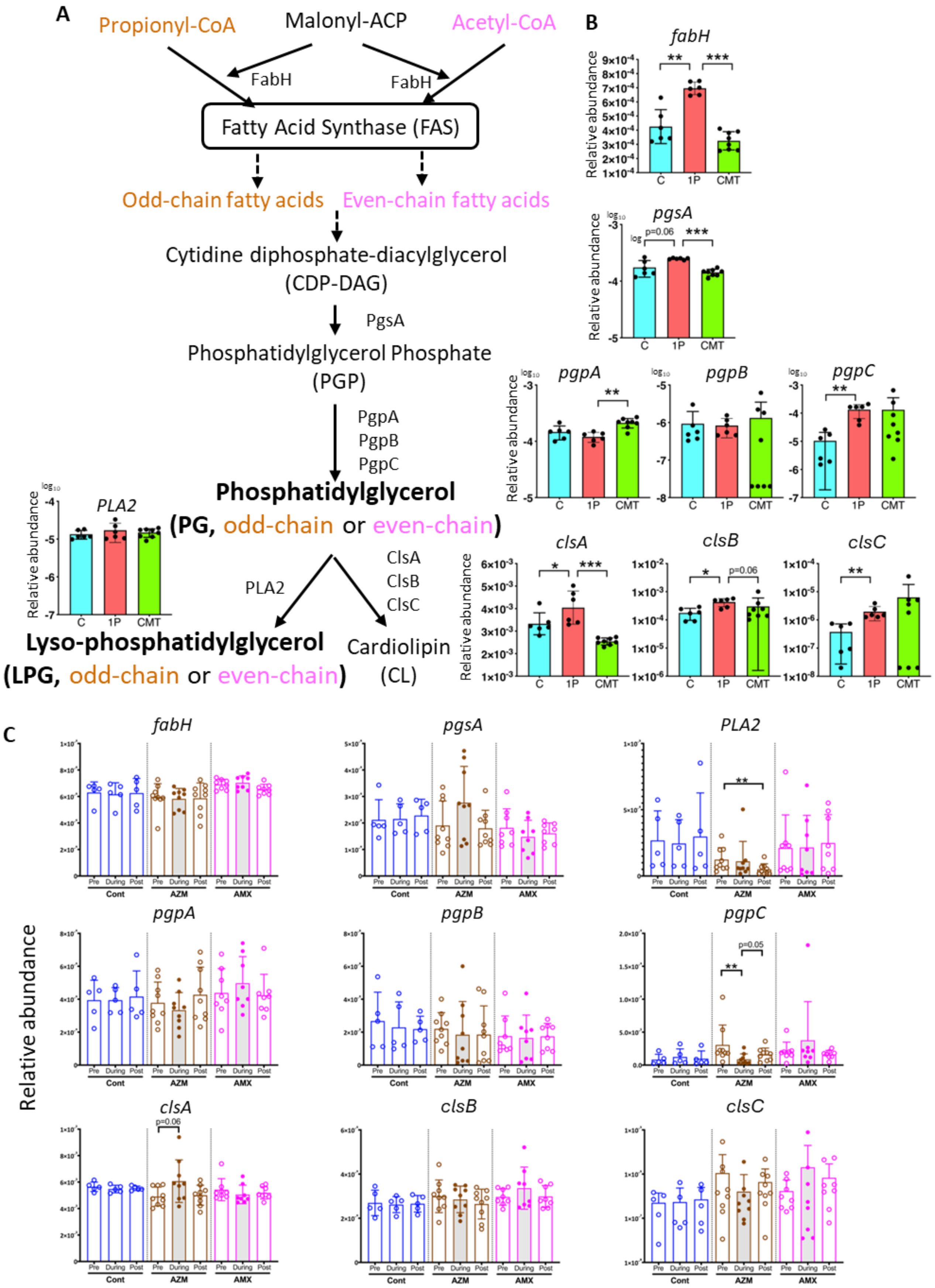
A
Propionyl-CoA
Malonyl-ACP
Acetyl-CoA
FabH
FabH
Fatty Acid Synthase (FAS)
Odd-chain fatty acids
Even-chain fatty acids
Cytidine diphosphate-diacylglycerol
(CDP-DAG)
PgsA
Phosphatidylglycerol Phosphate
(PGP)
PgpA
PgpB
PgpC
Phosphatidylglycerol
(PG, odd-chain or even-chain)
PLA2
ClsA
ClsB
ClsC
Lyso-phosphatidylglycerol
(LPG, odd-chain or even-chain)
Cardiolipin
(CL)
PLA2
B
fabH
pgsA
pgpA
pgpB
pgpC
clsA
clsB
clsC
Relative abundance
C
1P
CMT
p=0.06
C
fabH
pgsA
PLA2
pgpA
pgpB
pgpC
clsA
clsB
clsC
Relative abundance
Pre
During
Post
Cont
AZM
AMX
p=0.05
p=0.06

**Supplemental Figure 7. Key enzymes in the gut bacterial pathways of phosphatidylglycerol (PG) and lysyl-phosphatidylglycerol (LPG) biosynthesis. Panel A) Organization of the metabolic pathways.** The key enzymes include: FabH (β-ketoacyl-ACP synthase III), encoded by *fabH* ,which plays a critical role in initiating fatty acid synthesis and regulates the formation of odd-chain and even-chain fatty acid structures in PG and LPG; PgsA (phosphatidylglycerophosphate synthase), encoded by *pgsA*, which controls the crucial step in bacterial phospholipid biosynthesis, converting cytidine diphosphate-diacylglycerols (CDP-DAG) and glycerol-3-phosphate (G3P) into phosphatidylglycerophosphates (PGP); PgpABC (Phosphatidylglycerophosphatases A, B, C), encoded by *pgpA, pgpB, pgpC*, respectively, involved in the final step of bacterial PG biosynthesis from PGP (The functional redundancy of the 3 genes, all contributing to PG production enables bacteria to maintain a steady supply, which is crucial for membrane integrity); encoding phospholipase A2 (PLA2) that plays a critical role in phospholipid metabolism by hydrolyzing the sn-2 acyl bond of glycerophospholipids to generate lysophospholipids; ClsABC (Cardiolipin Synthases A, B, C), encoded by clsA, clsB, clsC, respectively, convert PG into cardiolipin (CL). **Panel B) Relative abundance of each gene based on metagenomic analysis of the cecal microbiome in NOD mice at P23.** Specimens altered by antibiotic exposure (1P) and restored by cecal material transplant (CMT), were compared with control (C). *p<0.05; **p<0.01; ***p<0.0001, compared with control by Mann-Whitney *U*-test. **Panel C) Relative abundance of each gene based on metagenomic analysis of the fecal microbiome of human subjects;** before (Pre), during, and after (post) antibiotic treatment. *p<0.05 compared with pre within the same group; Mann-Whitney *U*-test.

.

### Supplemental Tables

**Table S1.** Full list of the lipid compounds identified in lipidomic analysis of Experiments 1, 2 and 3. (Attached Excel file Table S1)

**Table S2.** The 391 compounds identified in lipidomic analysis of the 6 bacterial strains representative of the major intestinal bacterial taxa (Attached Excel file Table S2).

**Table S3.** Chemical characteristics of the 4 phospholipid compounds and their abundances in the Experiment 1 samples.

| Name | Chemical formula | Molecular weight | Fold-differences in abundance | | | |
|---|---|---|---|---|---|---|
| | | | $FD_{(Conv/GF)}$ (p-value)[a] | | $FD_{(Conv/Diet)}$ (p-value)[b] | |
| LPG(13:0) | $C_{19}H_{38}O_9P(Na)$ | 464.5 | 1233 | (p=0.002) | 192 | (p=0.016) |
| LPG(16:0) | $C_{22}H_{44}O_9P(Na)$ | 506.5 | 122 | (p=0.015) | 5.6 | (p=0.106) |
| LPG(18:0) | $C_{24}H_{48}O_9P(Na)$ | 534.6 | 126 | (p=0.089) | 22 | (p=0.212) |
| PG(15:0_15:0) | $C_{36}H_{70}O_{10}P(Na)$ | 716.9 | 1031 | (p=0.0003) | 34 | (p=0.004) |
| Retinoic acid[c] | $C_{20}H_{28}O_2$ | 300.44 | 1.4 | (p=0.527) | 1.0 | (p=0.983) |

[a.] Fold-difference between abundances in specimens from conventional and germ-free mice.
[b.] Fold-difference between abundances in specimens from conventional mice and diet samples.
[c.] Retinoic acid serves as a control lipid compound.

**Table S4.** List of primer sequences used for RT-qPCR analysis.

| Target | Forward primer | Reverse primer | Source |
|---|---|---|---|
| Mus musculus | | | |
| 18S | CATTCGAACGTCTGCCCTAT | CCTCCAATGGATCCTCGTTA | Zhang et al. 2021[1] |
| GAPDH | TGGTGAAGGTCGGTGTGAAC | CCATGTAGTTGAGGTCAATGAAGG | Lin et al. 2019[2] |
| NFκB1 | ATGGCAGACGATGATCCCTAC | CGGAATCGAAATCCCCTCTGTT | Wang et al. 2013[3] |
| RORγT | CCGCTGAGAGGGCTTCAC | TGCAGGAGTAGGCCACATTACA | Wang et al. 2013[3] |
| PPARγ | GTGATGGAAGACCACTCGCATT | CCATGAGGGAGTTAGAAGGTTC | Wang et al. 2013[3] |
| TLR5 | TGGGGACCCAGTATGCTAACT | CCACAGGAAAACAGCCGAAGT | Wang et al. 2013[3] |
| CASP1 | ACAAGGCACGGGACCTATG | TCCCAGTCAGTCCTGGAAATG | Wang et al. 2013[3] |
| NOS2 | GTTCTCAGCCCAACAATACAAGA | GTGGACGGGTCGATGTCAC | Wang et al. 2013[3] |
| REG3γ | ATGCTTCCCCGTATAACCATCA | ACTTCACCTTGCACCTGAGAA | Wang et al. 2013[3] |
| MUC2 | AGGGCTCGGAACTCCAGAAA | CCAGGGAATCGGTAGACATCG | Wang et al. 2013[3] |
| IL12 | CAATCACGCTACCTCCTCTTTT | CAGCAGTGCAGGAATAATGTTTC | Wang et al. 2013[3] |
| ECAM | CTGGCGTCTAAATGCTTGGC | CCTTGTCGGTTCTTCGGACTC | Wang et al. 2013[3] |
| ICAM1 | GTGATGCTCAGGTATCCATCCA | CACAGTTCTCAAAGCACAGCG | Wang et al. 2013[3] |
| Homo sapiens | | | |
| 18S | GTAACCCGTTGAACCCCATT | CCATCCAATCGGTAGTAGCG | Rho et al. 2010[4] |
| GAPDH | TGCACCACCAACTGCTTA | GGATGCAGGGATGATGTTC | Rho et al. 2010[4] |
| NFκB1 | GTCAAAAACGCCACCTCTCAA | CTCGCATGGAATTTGGAACCG | Wang et al. 2013[3] |
| RORγT | GTGGGGACAAGTCGTCTGG | AGTGCTGGCATCGGTTTCG | Wang et al. 2013[3] |
| IL13 | CCTCATGGCGCTTTTGTTGAC | TCTGGTTCTGGGTGATGTTGA | Wang et al. 2013[3] |
| TLR5 | GCCGGTCCTGTGTTTGGAAT | GGTGAGGTTGCAGAAACGATAAA | Wang et al. 2013[3] |
| TNFα | GGGTTGGGCAACAAGTATGTC | GGTGTCATCTCGGAGGTAATTCA | Wang et al. 2013[3] |
| CASP1 | TTTCCGCAAGGTTCGATTTTCA | GGCATCTGCGCTCTACCATC | Wang et al. 2013[3] |
